# Intrinsic self-organization of integrin nanoclusters within focal adhesions is required for cellular mechanotransduction

**DOI:** 10.1101/2023.11.20.567975

**Authors:** Kashish Jain, Kyna Yu En Lim, Michael P. Sheetz, Pakorn Kanchanawong, Rishita Changede

## Abstract

Upon interaction with the extracellular matrix, the integrin receptors form nanoclusters as a first biochemical response to ligand binding. Here, we uncover a critical biodesign principle where these nanoclusters are spatially self-organized, facilitating effective mechanotransduction. Mouse Embryonic Fibroblasts (MEFs) with integrin β3 nanoclusters organized themselves with an intercluster distance of ∼550 nm on uniformly coated fibronectin substrates, leading to larger focal adhesions. We determined that this spatial organization was driven by cell-intrinsic factors since there was no pre-existing pattern on the substrates. Altering this spatial organization using cyclo-RGD functionalized Titanium nanodiscs (of 100 nm, corroborating to the integrin nanocluster size) spaced at intervals of 300 nm (almost half), 600 nm (normal) or 1000 nm (almost double) resulted in abrogation in mechanotransduction, indicating that a new parameter i.e., an optimal intercluster distance is necessary for downstream function. Overexpression of α-actinin, which induces a kink in the integrin tail, disrupted the establishment of the optimal intercluster distance, while simultaneous co-overexpression of talin head with α-actinin rescued it, indicating a concentration-dependent competition, and that cytoplasmic activation of integrin by talin head is required for the optimal intercluster organization. Additionally, talin head-mediated recruitment of FHOD1 that facilitates local actin polymerization at nanoclusters, and actomyosin contractility were also crucial for establishing the optimal intercluster distance and a robust mechanotransduction response. These findings demonstrate that cell-intrinsic machinery plays a vital role in organizing integrin receptor nanoclusters within focal adhesions, encoding essential information for downstream mechanotransduction signalling.

## INTRODUCTION

Transmembrane signalling receptors such as integrins, cadherins, T-cell receptors, receptor tyrosine kinases, insulin receptors, and immunological synapses organize in nanoclusters which act as signalling hubs required for downstream signalling in growth and development^1–9^. There is some evidence that the nanoclusters of transmembrane signalling receptors mentioned above are self-organized in space^3,4, 9^, however, whether this spatial organization is required to bring about functional consequences is still an open question. We explore this question using integrin nanoclusters as an example of signalling hubs of transmembrane receptors that bind to extracellular ligands, such as cyclo RGD^3,10^. These integrin nanoclusters sense the physicochemical properties of the ECM (force geometry, growth factor signalling) ^1,3,11^, and bring about mechanotransduction which results in functional consequences in the short time scale where cell spread, actin organizes into stress fibres, YAP translocated to the nucleus^12, 13^ which intern direct long term functional consequence, for example, cell differentiation, and when it goes awry it can lead to devastating diseases such as cancer, and fibrosis^1, 2, 14–17^. Hence, it is important to understand what are the key biodesign parameters that govern this vital signalling process. One such crucial yet under-explored parameter could be the spatial organization of the signalling receptor clusters. Empirical evidence suggests that spacing between integrin nanoclustered subunits could be of the order of ∼500 nm^11, 17–19^. For example, fibroblasts spread on micropillars with 500 nm spacing appear to form continuous focal adhesion-like structures, whereas larger inter-pillar distance (>=1 micron) result in disjointed, rather than continuous, adhesion plaques^18–20^. In this manuscript, we address whether the spatial organization of integrin nanoclusters is precise thereby following an inherent biodesign principle and if so does this biodesign have functional consequences on mechanosignalling and mechanotransduction.

To probe biodesign principles of integrin spatial organization, nanopatterning offers an excellent way to specifically perturb the spatial organization of the nanoclusters by guiding them to form at predefined locations without altering any other biochemical parameter such as ligand inputs or signalling events in the extracellular or intracellular space. We nanopattern titanium and use these patterns to present cyclo-RGD, an integrin-binding ligand to fibroblast cells^11, 21^. Compared to more commonly used gold substrates, our method to functionalize titanium nanopatterns provides precise and quick functionalization, seamless passivation, and effective microscopic imaging with minimal fluorescence quenching^11,21,22^. In addition, Titanium functionalization can be further used to immobilize other ligands to investigate the impact of altered spatial organization on cellular functions.

The signals are probed by the cells and transmitted intracellularly to bring about function. Mechanosignalling and mechanotransduction recruit a multitude of proteins to the adhesion site that bind to the integrin β cytoplasmic tail^2, 23–26^. However, due to steric constraints, exchange, competition, and cooperation of integrin partners likely comprise major aspects of integrin regulatory mechanisms^26^. For instance, during early phase of nascent adhesion formation, α-actinin associates with putatively inactive integrins prior to the recruitment of paxillin and talin^27, 28^, before switching to associating with F-actin in stress fibers in mature adhesion^29^. Likewise, Talin has been shown to be recruited rapidly to nascent adhesions and plays multiple roles throughout the course of adhesion maturation. These include binding to integrins via the head and rod domains^2, 30^, promoting integrin activation, integrin clustering, talin head mediated recruitment of formin FHOD1 which polymerizes actin locally at the nascent adhesions, binding to cortical actin via the rod domain, stretching to bear load and tension, among others ^3, 30, 31^. The interplay between these adhesion proteins can help cells optimally respond to ECM physical cues such as force (substrate rigidity) and geometry ^32, 33^.

Using titanium nanopatterning^21, 34^, in this manuscript, we uncover an important biodesign principle of self-organization that is necessary for signalling downstream of activated transmembrane receptors using integrin receptors. In fibroblasts, for effective mechanosignal transduction, signalling hubs of integrin receptors nanoclusters have to be spaced with a precise intercluster distance of ∼550 nm between them. If this distance is halved or doubled, mechanosignal transduction does not proceed. This is driven by cell-intrinsic activation of the integrin cytoplasmic tail – talin – FHOD1 – actomyosin contractility at the integrin nanoclusters.

## RESULTS

### Integrin inter-cluster spacing regulates cell spreading, adhesion maturation and mechanosignalling

To understand whether integrin nanoclusters are randomly or precisely organized in space, we sought to characterize the spatial organization of integrin nanoclusters in focal adhesions. Integrin β3 depleted MEFs (using RNAi)^3^ expressing mEos2-Integrin β3 were allowed to spread on cyclo-RGD functionalized glass substrates for 15 min. Subsequently, they were fixed and imaged using Photo-activated Localization Microscopy (PALM) and the super resolution images were reconstructed using Picasso software^35, 36^. Integrin nanoclusters, formed upon ligand binding^3^, were analysed using DBSCAN cluster analysis. We observed that within focal adhesions, integrin nanoclusters were arranged with an average centre-to-centre distance of 555.16±204.23 (mean±s.d.; Fig. 1a). Since in these experiments, the ECM ligand was presented in a continuous manner with no apparent spacing, this inter-cluster spacing should be determined by a cell-intrinsic mechanism.

**Figure 1.**
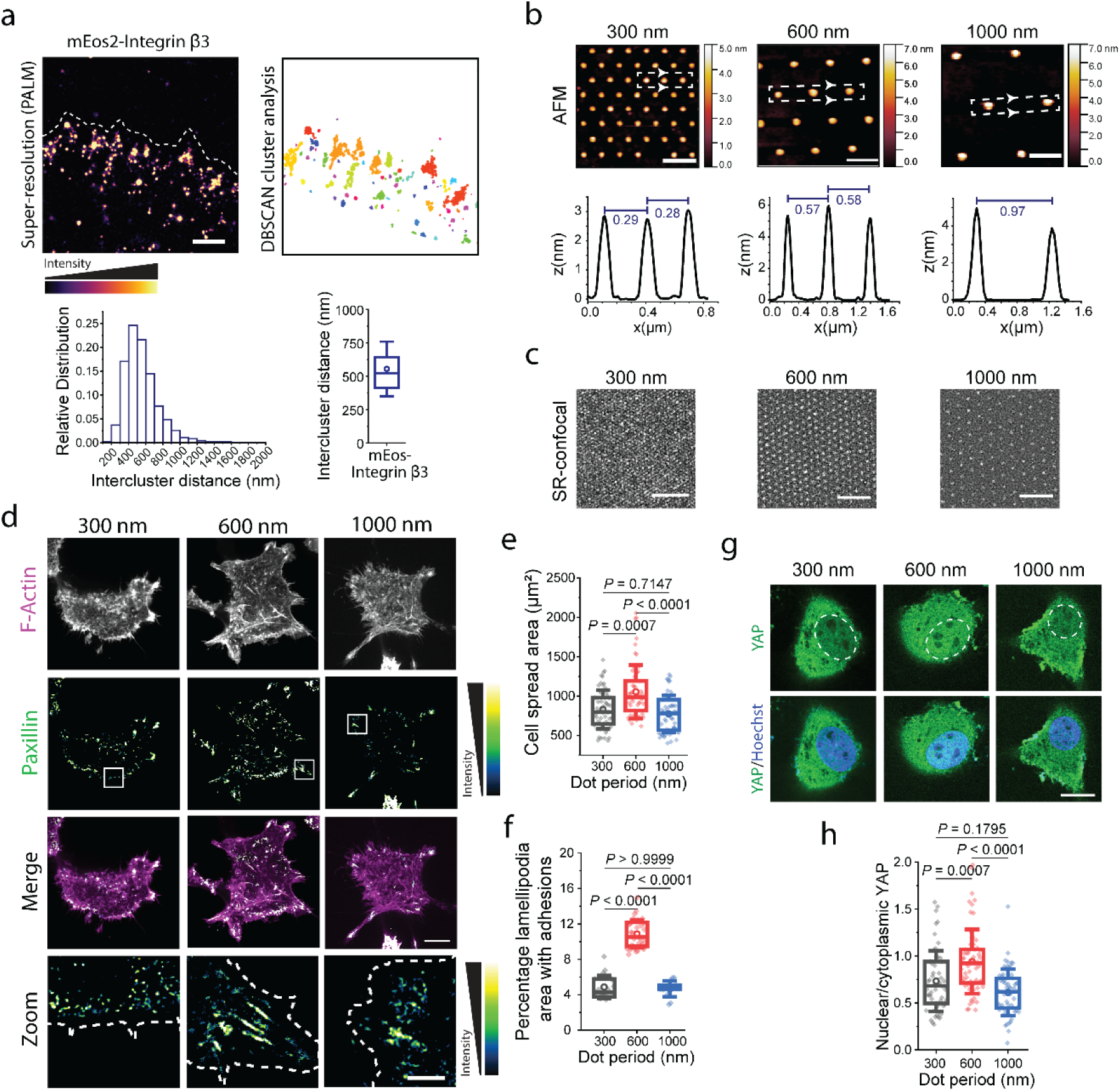
Optimal inter-cluster spacing of 600 nm promotes cell spreading, adhesion maturation and YAP nuclear translocation. **a**, Reconstructed PALM super-resolution image (left) of integrin β3 in nascent adhesions spread on fibronectin for 10-15 mins, and quantification of the integrin intercluster distance histogram (middle), and box plot (right). Scale bar, 500 nm. n=1702 from 6 cells. **b**, Representative super-resolution (SR) confocal image (left) of MEFs overexpressing integrin β3-GFP spread for 1 hr, along with zoom of white box (right). Relative frequency of the intercluster distance of integrin β3 nanoclusters. Scale bar (left) 10 µm. Scale bar (right) 2µm. **c**, Representative AFM images of the nanopatterned substrates, and linescans (middle) of the marked regions in the AFM images reporting the inter-disc distance. Scale bars, 500 nm. **d**, Confocal fluorescence images of the nanodisc substrates labelled with Dylight-405 labeled Neutravidin. Scale bars, 500 nm. **e**, Images of MEFs expressing EGFP-Paxillin with phalloidin staining for F-actin, along with the merge and zoom of the white box, spread on nanopatterns with different periods. Scale bar for Merge, 10 μm. Scale bar for zoomed inset, 3 μm. **f**, Quantifications of cell spread area (n ≥ 57 cells from three independent experiments, analyzed by the Kruskal–Wallis one-way ANOVA and Dunn’s post hoc test; *P* values are indicated in the figure) and **g**, percentage lamellipodia area with adhesions (n ≥ 14 cells from three independent experiments, analyzed by the Kruskal–Wallis test and Dunn’s post hoc test; *P* values are indicated in the figure). **h**, Confocal images showing YAP localization as a function of nanopattern periodicity. Nuclear boundary as marked by Hoechst staining indicated by whited dashed lines. Scale bar, 20 μm. **i**, Quantification of the nuclear to cytoplasmic YAP ratio (n ≥ 58 cells from three independent experiments, analyzed by the Kruskal–Wallis test and Dunn’s post hoc test; *P* values are indicated in the figure). Box plots with data overlay display the upper and lower quartiles and a median, the circle represents the mean, and the whiskers denote the standard deviation values, along with the individual points and P values. Detailed statistics are summarized in Supplementary Table 1.

To determine whether the observed inter-cluster spacing of ∼550 nm represents an important parameter for adhesion maturation or other cellular processes, we sought to present cells with a predefined spacing between receptor nanoclusters using nanopatterning. We presented cyclo-RGD ligand on titanium (Ti) nanodiscs of 100 nm diameter to resemble the diameter of integrin nanoclusters. These discs were arranged in hexagonal arrays with periodicity of 300, 600, or 1000 nm (Fig. 1b) to mimic half, normal, or double spacing, compared to observed spacing on continuous substrates. Characterization by Atomic Force Microscopy (AFM) verified the spacing and diameter of the nanodiscs used (Fig. 1b). To present cyclo-RGD to cells, the Ti discs were functionalized with biotinylated PIP_2_ lipids. To allow imaging of these substrates, Dylight-labelled Neutravidin was sandwiched between the biotinylated lipids and biotinylated cyclo-RGD^21^. The surrounding glass was passivated using supported lipid bilayers^21^. Imaging by confocal or structured-illumination-based super-resolution confocal microscopy similarly revealed the nanodisc pattern to conform closely to the expected periodicity (Fig. 1c). Subsequently, MEFs were spread on the cyclo-RGD functionalized nanodiscs for 1 h. Here, we observed that MEFs spread most favourably on the 600 nm nanopatterns, exhibiting a significantly larger cell area (1055.7±44.67 µm^2^, mean ± s.e.m.) compared to cells spread on 300 nm (830.4±31.3 µm^2^, mean ± s.e.m.) and 1000 nm patterns (773.1±31.5 µm^2^, mean ± s.e.m.; Fig. 1d,e). We next quantified the extent of focal adhesion formation, defined as the sum of the focal adhesion area normalized to the lamellipodia area^11^. We again observed maximum extent of focal adhesion formation on 600 nm nanopattern, and it was ∼2.3 fold higher compared to on either 300 nm or 1000 nm nanopattern (Fig. 1d,f). In conjunction with this, we noted that MEFs spread on the 600 nm nanodisc period formed longer adhesions with a higher adhesion area (Fig. 1d, Fig. S1), while focal adhesions formed on the 300 nm or 1000 nm nanodiscs were smaller and had a lower adhesion area (Fig. 1d, Fig. S1). This shows that there is an inherent biodesign that brings about required spatial organization of the integrin nanocluster spacing which is necessary for focal adhesion formation, maturation, and cell spreading. This biodesign principle governs that ∼600 nm is an optimal intercluster spacing, while ∼300 nm (half) or ∼1000 nm (double) were suboptimal to bring about effective mechanotransduction.

Next, we assessed the effect of nanopattern periodicity on downstream hallmarks of mechanosignalling and mechanotransduction i.e. increase in FAK phosphorylation and translocation of YAP to the nucleus as measured by nuclear to cytoplasmic ratio of GFP-YAP (similar to experiments reported in ^12, 13^). We observed a ∼2 times higher FAK phosphorylation on 600 nm compared to either 300 nm or 1000 nm spacing (Fig. S2a,b), suggesting again that 600 nm acted as the optimal biodesigned spacing between nanoclusters, while 300 nm or 1000 nm were suboptimal. Similarly, when we measured the nuclear localization of YAP as normalized to cytoplasmic YAP, we observed a higher nuclear to cytoplasmic ratio in cells spread on 600 nm (0.94±0.04, mean±s.e.m.) compared to 300 nm (0.73±0.04, mean±s.e.m.) or 1000 nm spacing (0.61±0.03, mean±s.e.m.), indicative of the significantly higher mechanotransduction on the 600 nm spaced nanopatterns (Fig. 1g,h). Taken together, our results showed that cell spreading, extent of focal adhesion formation, and downstream mechanosignalling are strongly determined by the inter-cluster spacing between integrin nanoclusters, with the optimal nanocluster spacing of ∼600 nm.

### Overexpression of the integrin binding domain (SR12) of α-actinin 1 disrupts establishment of integrin inter-cluster spacing

Next, we sought to delineate the molecular mechanism that establishes this precise spatial organization of integrin nanoclusters required for adhesion maturation and mechanotransduction. We conducted a mini overexpression screen to identify which focal adhesion components could govern this spatial organization. We observed that when α-actinin 1 was overexpressed in MEFs, cells spread to a similar extent on all three inter-cluster spacing substrates (2358.91±194.03 µm^2^ on 300 nm, 2238.40±171.91 µm^2^ on 600 nm, 2476.17±195.94 µm^2^ on 1000 nm; mean±s.e.m.), and the higher spreading on the optimal 600 nm spacing is lost (P=0.89 for 300 nm vs 600 nm, P=0.69 for 600 nm vs 1000 nm; Fig. 2a,b). Likewise, the focal adhesion area was comparable for cells on substrates with all 3 spacings (0.56±0.04 µm^2^ on 300 nm, 0.58±0.01 µm^2^ on 600 nm, 0.57±0.02 µm^2^ on 1000 nm; mean±s.e.m.). Focal adhesions majorly formed in the cell interior and were thin, long and segmented, as compared to normal FAs which are thick and continuous (comparing Fig. 2a and Fig. 1d). In addition, overexpression of α-actinin 4, another mammalian α-actinin isoform ubiquitously expressed in non-muscle cells^37^, resulted in a similar loss of establishment of optimal inter-cluster spacing (P=0.34 for 300 nm vs 600 nm, P=0.23 for 600 nm vs 1000 nm; Fig. S3a,b). Since α-actinin 1 could bind to both integrin and actin as well as other adhesion-associated proteins, we next sought to identify the specific functional modules responsible for disrupting inter-cluster spacing establishment. We found that the overexpression of the integrin-binding domain of α-actinin 1 which encompasses the first and second spectrin repeat (called SR12 hereafter)^38,39^, resulted in the loss of establishment of optimal integrin inter-cluster spacing, as indicated by comparable cell spreading area for all three nanocluster spacing (P>0.9999 for 300 nm vs 600 nm, P>0.9999 for 600 nm vs 1000 nm; Fig. 2c,d). Since α-actinin also functions as F-actin crosslinkers, we tested whether the specific integrin binding domain or the actin crosslinking activity of α-actinin 1 may contribute to the loss of appropriate establishment of integrin inter-cluster spacing. We tested this supposition by overexpressing of another actin crosslinked, viz, the N93K mutant of Myosin IIA that is known to only crosslink actin and has no ATPase or motor activity^27, 40^. We observed that cells overexpressing Myosin IIA-N93K still exhibited a higher cell spreading on the optimal 600 nm intercluster distance compared to 300 nm or 1000 nm (P=0.0011 for 300 nm vs 600 nm, P=0.0017 for 600 nm vs 1000 nm) similar to control MEFs (Fig. 2e,f). These results suggest that any additional actin crosslinking function due to overexpression of α-actinin 1 did not contribute to the establishment of the optimal integrin inter-cluster spacing. Taken together, our results identified that the integrin-binding domain of α-actinin inhibited the ability of cells to establish the optimal 600 nm integrin inter-cluster spacing.

**Figure 2.**
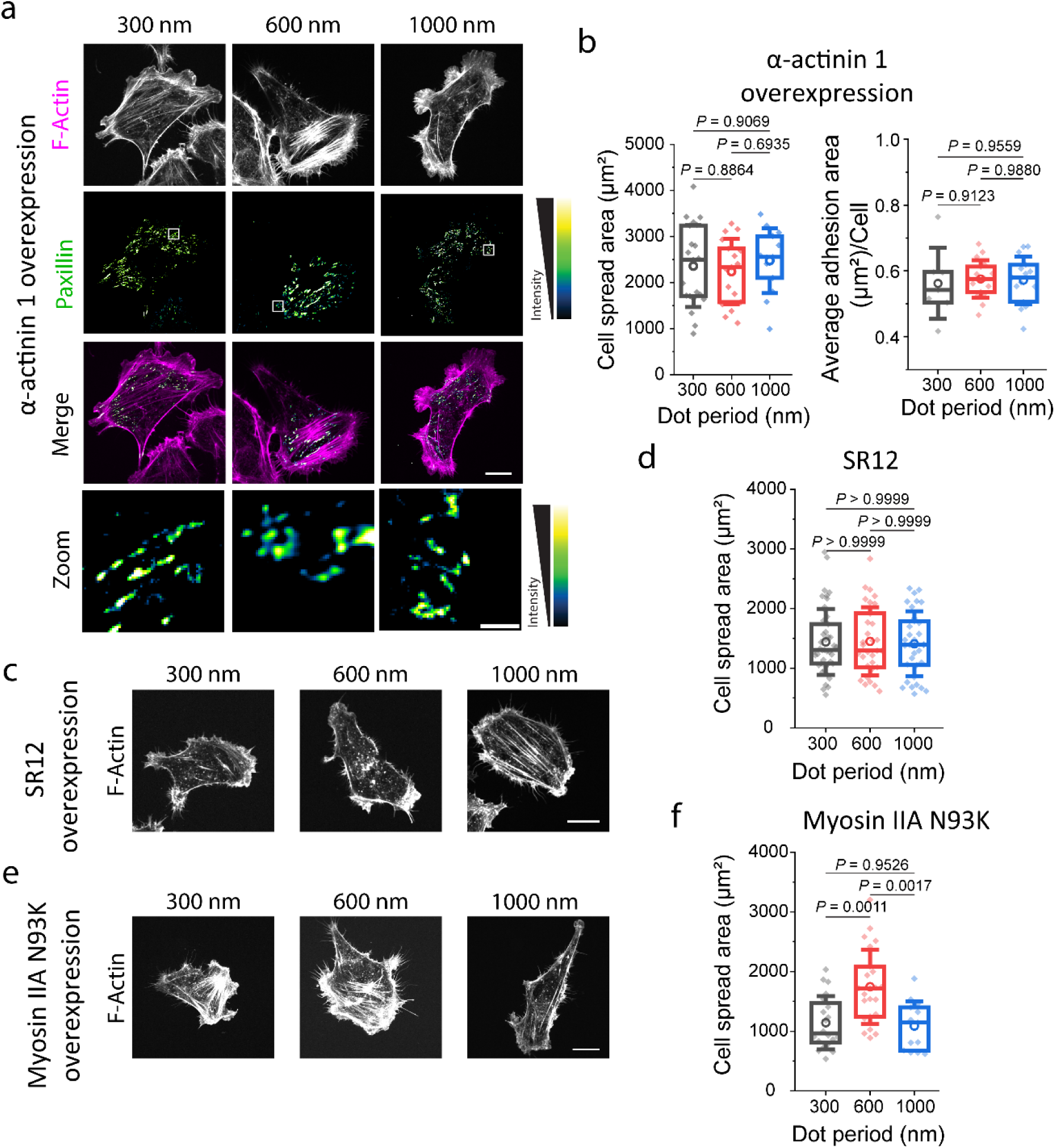
Overexpression of α-actinin 1 integrin binding domain (SR12) disrupts integrin inter-cluster spacing establishment. **a**, F-actin and paxillin immunofluorescence images of MEFs overexpressing mCherry-α-actinin 1 spread on different nanodisc periods, along with the merge and zoom of the white box in paxillin. Scale bar for merge, 15 µm. Scale bar for zoom, 2 µm. **b**, Quantifications of cell spread area (n ≥ 13 cells from three independent experiments, analyzed by one-way ANOVA and Tukey’s post hoc test; *P* values indicated in the figure) and average adhesion area/cell (n ≥ 14 cells from three independent experiments, analyzed by Kruskal–Wallis one-way ANOVA and Dunn’s post hoc test*; P* values indicated in the figure). **c**, Representative F-actin images of cells overexpressing EGFP-SR12 on different disc periods along with **d**, quantification of cell spread area (n ≥ 31 cells from three independent experiments, analyzed by Kruskal– Wallis test and Dunn’s post hoc test*; P* values indicated in the figure). Scale bar, 15 µm. **e**, Representative F-actin images of cells overexpressing mCherry-Myosin IIA-N93K mutant on different disc periods along with **f**, quantification of cell spread area (n ≥ 13 cells from three independent experiments, analyzed by one-way ANOVA and Tukey’s post hoc test*; P* values indicated in the figure). Scale bar, 15 µm. Box plots with data overlay display upper and lower quartiles and median, circle represents the mean, and whiskers denote the standard deviation, along with individual data points. Detailed statistics are summarized in Supplementary Table 2.

### Interaction between integrin β3 and α-actinin SR12 disrupts establishment of integrin inter-cluster spacing and cell spreading

The integrin-binding SR12 domain of α-actinin binds to integrin β cytoplasmic tails at the cytoplasmic domain in close proximity to the talin head binding site^26, 41^. The SR12 domain can compete with the talin head for the integrin β3 tail, whereas it can bind in a cooperative manner with the talin head domain to the β1 tail^39^. To identify which β integrin mediates our observed phenotypes, we perturbed specific integrin binding to the substrate by G-pen peptide that blocks integrin β3 or HMB1-1 antibody that blocks β1 integrin. We observed that blocking the integrin β3 subunit inhibited the establishment of optimal inter-cluster spacing (P=0.91 for 300 nm vs 600 nm, P=0.98 for 600 nm vs 1000 nm; Fig. 3a,b) whereas, we observed a significantly higher cell spreading on 600 nm nanopatterns as compared to 1000 nm with integrin β1 blocking (P<0.0001 for 600 nm vs 1000 nm; Fig. 3a,c). Thus, the binding activity of integrin β3, but not integrin β1, is required for establishing the cell-intrinsic biodesign principle for optimal spatial organization of integrin nanoclusters.

**Figure 3.**
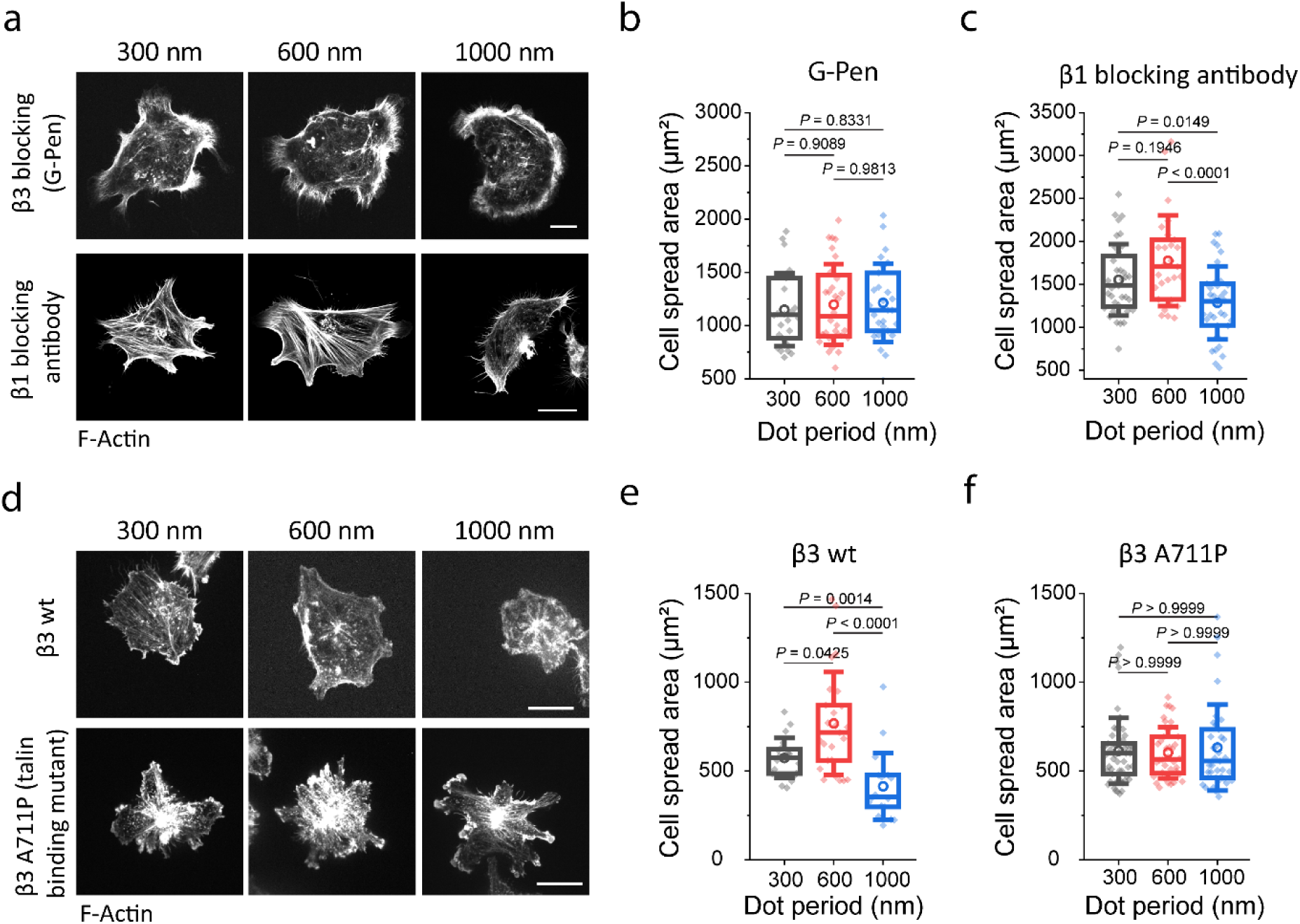
Integrin β3 but not integrin β1, through its talin binding function, is involved in establishing inter-cluster spacing. **a**, Representative images of cells spread on the 300, 600, and 1000 nm disc diameter patterns treated with the β3 blocking peptide-G-Pen, or the β1 blocking antibody, HMβ1-1 clone, stained for F-actin. Scale bar (top), 10 µm. Scale bar (lower), 20 µm. **b**, Quantification of cell spread area (n ≥ 26 cells from 3 independent experiments, analyzed by one-way ANOVA test and Tukey’s post hoc test) for cells treated with G-Pen. **c**, Quantification of cell spread area (n ≥ 27 cells from three independent experiments, analyzed by one-way ANOVA test and Tukey’s post hoc test) for cells treated with β1 blocking antibody. **d**, Representative F-actin images of CHO cells overexpressing integrin β3 wt or β3 A711P mutant on different disc periods. Scale bar (top), 15 µm. Scale bar (bottom), 15 µm. Quantifications of cell spread area for **e**, β3-wt (n ≥ 19 cells from three independent experiments, analyzed by one-way ANOVA and Tukey’s post hoc test) and **f**, β3-A711P (n ≥ 36 cells from three independent experiments, analyzed by Kruskal-Wallis test). Box plots with data overlay display upper and lower quartiles and median, circle represents the mean, and whiskers denote standard deviation, along with individual data points. Detailed statistics are summarized in Supplementary Table 3.

Since SR12 could potentially compete with talin for integrin β3 binding, we hypothesized that SR12 overexpression might reduce integrin β3-talin interaction, and force transduction, thereby obstructing adhesion maturation. We focused on adhesion formation during early spreading on fibronectin since it predominantly involves integrin β3 rather than integrin β1^3,11,39,42^, consistent with our earlier observation that β3 inhibition abrogates the establishment of optimal inter-cluster spacing above. We observe that when MEFs overexpressing SR12 were plated for 15-20 mins on fibronectin-coated substrates, a significant reduction in cell spread area by 17.5% was observed (P<0.0001; Fig. S4a, b). Furthermore, the extent of adhesion maturation as measured by the total focal adhesion area normalized to the cell area (P<0.0001; Fig. S4a,c) was reduced by 30%. Similarly, marked decreases were also observed for both adhesion area (P<0.0001; Fig. S4d) and adhesion length (P<0.0001; Fig. S4e). However, after 1 hr spreading, we observed a 37% increase in cell spreading with SR12 overexpression (P<0.0001; Fig. S5a,c). This increase might be due to the differential interaction of SR12 with integrin β3 and β1, since integrin β1 is recruited after 1 hr spreading^39^. This can also explain the increase in cell spreading on the nanodisc substrates with α-actinin 1 overexpression, even without adhesion maturation.

Since the SR12 domains of the 1- and 4- isoforms of α-actinins showed an 88% similarity according to NCBI BLAST (Fig. S3c), we also evaluated the structural similarities between both the isoforms. We obtained r.m.s.d. value of 0.49 Å and a TM-score of 0.9925 (Fig. S3d), indicative of a highly conserved structure, suggesting that α-actinin-4 likely also binds to the integrin β3 and exerts similar phenotype to α-actinin 1, consistent with our empirical observation above (Fig. S3a,b).

Molecular dynamics simulations have earlier suggested that the presence of α-actinin may reduce the binding of integrin β3 to talin head domain^43^ by inducing a kink in the integrin β3 transmembrane domain and locking it in an inactive form. A point mutation A711P in integrin β3 could mimic this kink^43^, which inhibited integrin activation, and reduced HEK293FT cell spreading on PAC-1 and fibrinogen^44, 45^. To test this, we aimed to replace the native integrin with this point mutant and observe its effect on establishing inter-cluster spacing. For this purpose, we used CHO.K1 cells which have intrinsically low integrin expression^46^. We found that compared to the expression of wild-type β3, the expression of β3-A711P in CHO cells led to a 24% reduction of cell spread area (P=0.0003) and a 41% reduction in total adhesion area normalized to the cell area (P<0.0001; Fig. S6a,b). Consistent with this, when CHO cells expressing β3-wt or β3-A711P were spread on the nanopattern substrates, we observed no significant difference in cell spreading on different nanopattern were observed for β3-A711P expressing cells (P=0.04 for 300 nm vs 600 nm, P<0.0001 for 600 nm vs 1000 nm; Fig. 3d,f) as compared to β3-wt (P>0.9999 for 300 nm vs 600 nm, P>0.9999 for 600 nm vs 1000 nm; Fig. 3d,e). These observations thus indicate that the A711P mutation in integrin β3 was sufficient to disrupt the integrin inter-cluster spacing establishment. This also further implicates α-actinin 1 induced conformational change of integrin β3 tail is the underlying mechanism that abrogates the biodesign principle that establishes the optimal integrin inter-cluster spacing. Since the loss of sensing is due to a mutation in integrin β3 cytoplasmic tail, it further emphasizes the cell intrinsic nature of establishment of appropriate intercluster spacing that is required for downstream mechanotransduction.

### Competition between talin head domain and α-actinin SR12 regulates establishment of integrin inter-cluster spacing

Our results thus far suggest that the overexpression of α-actinin SR12 domain may serve to decrease talin binding to integrin β3 by inducing a β3 tail kink formation. Since SR12 overexpression was earlier shown not to affect the expression levels of talin or integrin^39^, our results imply a direct competition between α-actinin and talin for integrin β3 binding. This suggests that the ability of cells to establish inter-cluster spacing and promote adhesion maturation is expected to depend on the relative concentration of SR12 to talin. In other words, high concentration of SR12 can outcompete talin for integrin β3 binding. If so, then co-overexpression of talin and SR12 could restore the concentration balance of these two proteins, thereby rescuing adhesion maturation and proper establishment of the inter-cluster spacing. To test this, we next co-overexpressed full-length talin and SR12 in MEFs and quantified their adhesion formation on unpatterned glass substrate. To maintain similar overexpression levels, we used equal concentrations of both the plasmids and imaged cells expressing both plasmids only. We observed that compared to cells only expressing SR12, cells expressing both full-length talin and SR12 were able to significantly increase cell spreading (P=0.0017 for SR12 vs TFL), extent of adhesion formation (Fig. 4a,b,c), and the ratio of total adhesion area by cell area (P<0.0001 for SR12 vs TFL). Altogether, these observations show that counterbalancing concentrations of talin and SR12 can rescue adhesion maturation.

**Figure 4.**
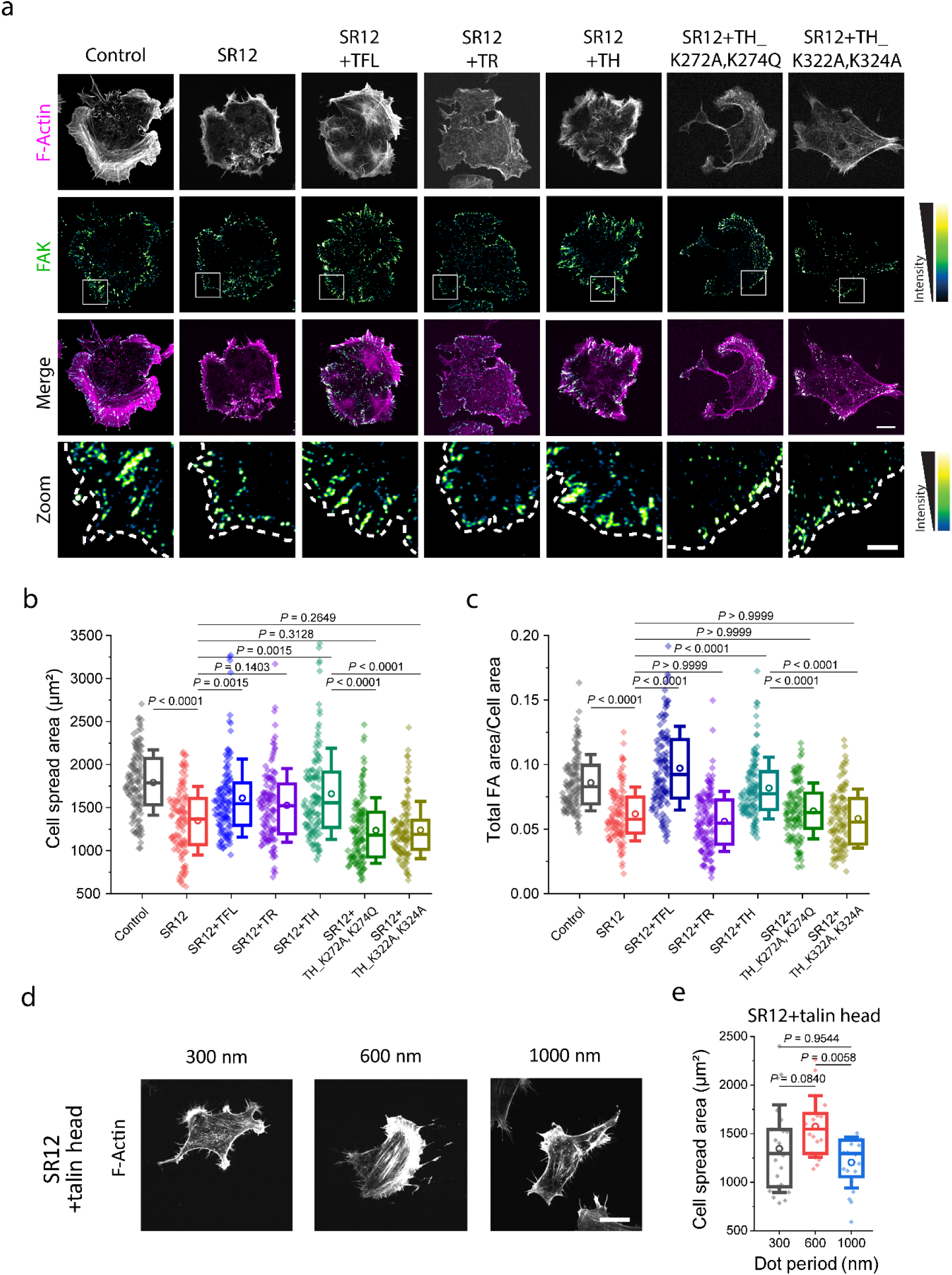
Cell establishment of inter-cluster spacing is fine-tuned by competition between talin head domain and α-actinin SR12 domain for integrin β3 binding. **a**, Control, SR12 overexpressing, SR12 and Talin full length (TFL) co-overexpressing, SR12 and Talin rod (TR) co-overexpressing, SR12 and Talin head (TH) co-overexpressing, SR12 and Talin head_K272A, K274Q co-overexpressing, SR12 and Talin head_K322A, K324A co-overexpressing cells were spread on fibronectin-coated glass dishes for 15-20 mins and immunostained for FAK and actin (phalloidin). Representative images showing F-actin, FAK, merge, and zoom of the white box in FAK. Scale bar for Merge, 10 µm. Scale bar for Zoom, 3 µm. **b**, Quantification of the cell spread area (n >= 107 cells from three independent experiments, analyzed by Kruskal-Wallis test and Dunn’s post hoc test; P values are indicated), **c**, total adhesion area normalized to cell area (analyzed by Kruskal-Wallis test and Dunn’s post hoc test; P values are indicated in the figure). **d**, Representative images of cells spread on the 300, 600, and 1000 nm disc diameter patterns co-overexpressing SR12 and talin head, stained for F-actin. Scale bar, 20 µm. **e**, Quantification of cell spread area (n ≥ 18 cells from three independent experiments, analyzed by Kruskal-Wallis test and Dunn’s post hoc test; P values are indicated). All graphs show individual points along with box plots of interquartile distance, line represents median, circle represents the mean, whiskers represent standard deviation. Detailed statistics are summarized in Supplementary Table 4.

Since the integrin binding sites in talin head and rod is far apart and the talin head binding site competes with SR12^39,43,47^, we next sought to rescue α-actinin SR12 phenotype with distinct talin domain co-overexpression. We co-overexpressed either the talin head domain (F0-F3, residue 1-433) or the talin rod domain (R1-DD, residue 453-2541) along with SR12 and spread MEF cells on fibronectin-coated glass substrates. We observed that co-expression with talin head domain but not talin rod domain is able to rescue the SR12 phenotype, as indicated by higher cell spreading, ratio of total adhesion area by cell area, and formation of larger adhesions (Fig. 4a,b,c). Further, to confirm the role of talin head in this biodesign, we made two distinct lysine point mutations at two locations each in the F2 and F3 domains of talin head (K272A, K274Q, and K322A, K324A) which has been shown to inhibit the interaction with PI(4,5)P_2_, disrupt integrin binding, and hinder adhesion maturation^48^. MEFs co-overexpressing either one of these two talin head double mutants (TH-K272A, K274Q, and TH-K322A, K324A) along with SR12 were observed to phenocopy SR12 overexpression cells, with a significant reduction in the cell spread area (P<0.0001 for both SR12 vs TH-K272A, K274Q and SR12 vs TH-K322A, K324A), and the ratio of total focal adhesion area by cell area (P<0.0001 for both SR12 vs TH-K272A, K274Q and SR12 vs TH-K322A, K324A) (Fig. 4a, b, c). Altogether these results provide empirical evidence that there is a concentration-dependent competition of SR12 with the integrin-binding domain of talin head and an appropriate balance of these two proteins is required for the proper adhesion maturation.

Hence, we hypothesized that co-overexpression of talin head domain along with SR12 can rescue the loss of inter-cluster spacing establishment by overexpression of SR12. We observed that SR12+talin head co-overexpression was able to demonstrate a significantly higher cell spreading when spread on 600 nm spacing as compared to 300 nm or 1000 nm spacing, suggesting the proper establishment of the integrin inter-cluster spacing (P=0.08 for 300 nm vs 600 nm, P=0.006 for 600 nm vs 1000 nm; Fig. 4d,e). This suggests that talin head is necessary to establish the optimal inter-cluster spacing and might be responsible for intrinsically organizing integrin nanoclusters at a distance of ∼600 nm. In addition, this demonstrates that tail rod domain-mediated attachment of integrin clusters to cortical actin is not responsible for the establishment of the spatial organization of integrin nanoclusters.

Since talin head is known as a key integrin activator in adhesion-dependent mechanotransduction^45, 49^, we next sought to determine whether activating integrins extrinsically may lead to organization of inter-cluster spacing. To test this, we treated the SR12 overexpressing cells with Mn^2+^, which is known to shift integrins into active conformation^50^. In SR12 overexpressing cells, unlike controls treated with Mn^2+^ we observed no significant difference upon Mn^2+^ treatment both in terms of the cell spread area (P>0.9999 for SR12 vs SR12+Mn^2+^; Fig. S7a,b), and total focal adhesion area normalized to the cell area (P>0.9999 for SR12 vs SR12+Mn^2+^; Fig. S7a,c). We thus conclude that integrin activation at extracellular domain, by Mn^2+^ cannot override the cell-intrinsic effect caused by SR12 overexpression on β3 cytoplasmic tail and kink free activation of cytoplasmic domain of integrin β3 tail is necessary for cell-intrinsic establishment of integrin inter-cluster distance.

### Local actin polymerization and actomyosin contractility are required for establishing integrin inter-cluster spacing

We then sought to investigate how talin head mediated the spatial organization of integrin inter-cluster distance. Talin head domain promotes recruitment of FHOD1 formin to nascent adhesions, which in turn promote local actin polymerization at integrin nanoclusters^3, 31^. Thus, local actomyosin contractility can play critical role in the establishment of the integrin inter-cluster distance. Therefore, we next sought to test the role of local actin polymerization by FHOD1 in establishing inter-cluster spacing. We knocked down FHOD1 using shRNA in cells that are co-expressing talin head and SR12. For cells spread on fibronectin-coated unpatterned substrates, we observed a significant reduction in cell spread area in the SR12+TH+FHOD1 shRNA group as compared to both control (P<0.0001 for Control vs SR12+TH+FHOD1 shRNA) and SR12+TH groups (P=0.0008 for SR12+TH vs SR12+TH+FHOD1 shRNA; Fig. 5a,b). We also observed a reduction in adhesion maturation, as shown by a significant decrease in total adhesion area normalized to cell area in SR12+TH+FHOD1 shRNA compared to both control (P=0.0001 for Control vs SR12+TH+FHOD1 shRNA) and SR12+TH groups (P=0.0001 for SR12+TH vs SR12+TH+FHOD1 shRNA; Fig. 5a,c). Furthermore, when FHOD1 shRNA expressing MEFs are plated on nanopattern substrates with different disc periods, we observed a loss of establishment of the inter-cluster spacing (P=0.98 for 300 nm vs 600 nm, P=0.95 for 600 nm vs 1000 nm; Fig. 5d,e). Our observations suggest that talin head is unable to rescue the SR12 phenotype in the absence of FHOD1. Since FHOD1 is known to promote anti-parallel actin polymerization at the adhesions^51^, we conclude that the establishment of inter-cluster spacing likely operate through talin head-mediated recruitment of FHOD1, whose local actin polymerization activity could then serve to define the optimal length scale for downstream mechanotransduction responses. The role of local actin polymerization cannot be studied directly as actin polymerization is requires for establishment of nanoclusters itself^3^, hence when no cluster formation is observed we were unable to study the establishment of proper intercluster distance.

**Figure 5.**
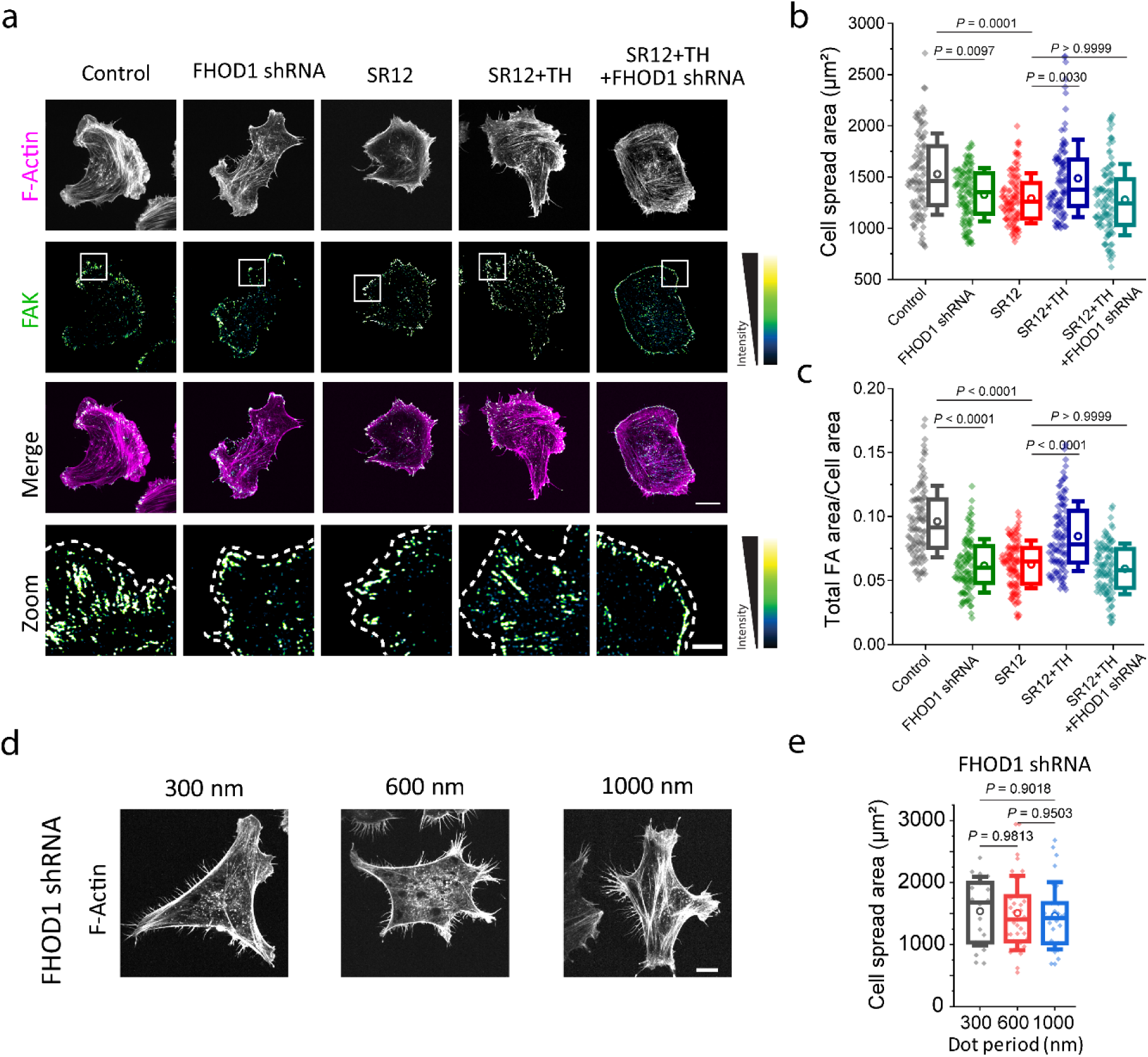
FHOD1 formin is required for establishing inter-cluster spacing and adhesion maturation. **a**, Control, FHOD1 shRNA expressing, SR12 expressing, SR12 and Talin head (TH) co-overexpressing, and SR12, TH and FHOD1 shRNA co-overexpressing cells were spread on fibronectin-coated glass dishes for 15-20 mins and immunostained for FAK and actin (phalloidin). Representative images showing F-actin, FAK, merge, and zoom of the white box in FAK. Scale bar for Merge, 10 µm. Scale bar for Zoom, 3 µm. **b**, Quantification of the cell spread area (n >= 97 cells from three independent experiments, analyzed by Kruskal-Wallis test and Dunn’s post hoc test; *P* values are indicated), **c**, total adhesion area normalized to the cell area (analyzed by Kruskal-Wallis test and Dunn’s post hoc test; *P* values are indicated). **d**, Representative images of cells spread on the 300, 600, and 1000 nm disc diameter patterns expressing FHOD1 shRNA, stained for F-actin. Scale bar, 20 µm. **e**, Quantification of the cell spread area (n ≥ 18 cells from three independent experiments, analyzed by One-way ANOVA test and Tukey’s post hoc test; *P* values are indicated). All graphs show individual points along with box plots of interquartile distance, the line represents the median, the circle represents the mean and the whiskers represent the standard deviation. Detailed statistics are summarized in Supplementary Table 5.

We further investigated if actomyosin contractility in the locally polymerized actin is required for the establishment of integrin inter-cluster spacing. To test whether myosin II contractility contributes to organization of the inter-cluster spacing, we inhibited myosin II activity using a low concentration (5 µM) of blebbistatin and quantified the extent of adhesion formation and maturation on the Ti nanodiscs. We observed that cell spread area on all three different nanocluster spacing (P=0.07 for 300 nm vs 600 nm, P=0.75 for 600 nm vs 1000 nm; Fig. S8b) were comparable. However, organized focal adhesions were not observed, but integrin nanoclusters formed on Ti discs (Fig. S8a). These results suggest a step-wise distinction between the myosin II-independent formation of nascent integrin nanoclusters, and the subsequent myosin II-dependent organization of integrin nanoclusters at a spacing of ∼600 nm. This also implicates establishment of inter-cluster spacing as a key mechanosensing checkpoint gating the maturation of nanoclusters into focal adhesions.

## DISCUSSION

Although the nanocluster organization of focal adhesions have been extensively documented by a wide range of super-resolution microscopy observation, the spatial organization of these nanoclusters, as well as their functional implications, has remained elusive and challenging to probe^15,25^. In this study, we show that the optimal inter-cluster distance between these nanoclusters is required for downstream intracellular signaling and function in cells. When we measured the inter-cluster distance between these nanoclusters, we observed a well-defined integrin nanocluster are intrinsically spaced with ∼550 nm intervals on uniformly coated ECM-coated substrates. A uniform substrate implies that this inter-cluster distance is established intracellularly and does not depend on external ligand presentation. In addition, we show that cytoplasmic tail domain of β3 integrin and not the extracellular domain of this transmembrane receptor is required to establish this inherent biodesign to spatially organize integrin nanoclusters.

Conversely, by making use of our recent Ti nanopatterning and functionalization method^21^ we were able to impose different inter-cluster spacing distances, we found that such spacing exerts profound effects on focal adhesion maturation and downstream mechanosignal transduction. Here, we focus on the 1 hr timepoint because it is sufficient to demonstrate effects on cell spreading and adhesion maturation, without the additional secretion of cell matrix proteins like collagen or fibronectin by cells. The downstream effects of integrin nanocluster spacing, dependent on FAK signalling, adhesion maturation and YAP nuclear translocation might occur at later timepoints. Effects of actin perturbations and talin mutants on unconstrained nascent adhesions have been demonstrated in our previous study^3^. Nanopatterned substrates provide us with the ability to perturb the intercluster distance, to look at downstream effects, which is not possible on continuous glass substrates. Nanodisc substrates also restrict integrin nanoclusters to be formed at a precise spatial distribution in space, as compared to uniformly coated substrates on which integrin nanoclusters can form at any distance, and are mobile^3^. We observed that cells require this biodesign principles to robustly self-organize the integrin inter-cluster spacing at an optimal distance of ∼550 nm, and exhibit a decreased mechanotransduction and signaling if the distance between these nanoclusters is physically halved- 300 nm, or almost doubled-1000 nm (Fig. 6) without changing any other biochemical or physical parameter presented to the cells. Cells on all three patters are within the same media on the same coverslip as all patters are patterned side by side on the same coverslip. The establishment of optimal integrin inter-cluster spacing described here shows that along with biochemical input from receptor-ligand binding, a physical parameter of receptor nanocluster spacing is necessary for adhesion maturation and proper mechanotransduction. There is a cell intrinsic biodesign established by which a precise inter-cluster distance of ∼550 nm is optimal between integrin nanoclusters.

**Figure 6.**
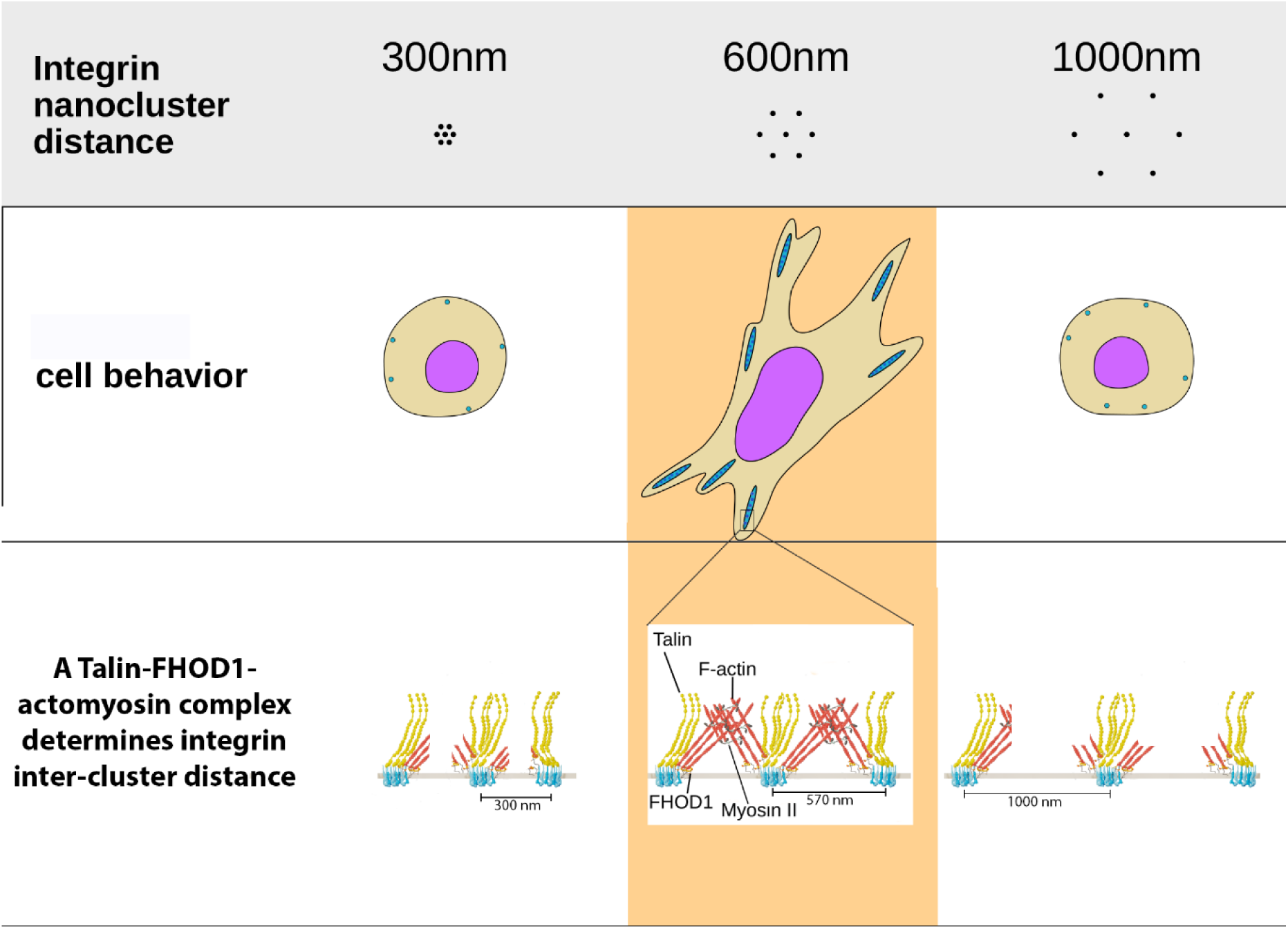
Proposed model for how integrin nanocluster distance regulate cell spreading and adhesion maturation. Cells establish an optimal 600 nm inter-cluster spacing and show a decrease in cell spreading, adhesion formation, and YAP signalling when inter-cluster spacing is changed to 300 nm or 1000 nm. Establishment of inter-cluster spacing depends on FHOD1 formin recruited to the integrin clusters in the presence of talin head domain, to promote local actin polymerization. Conversely, FHOD1 depletion or displacement of talin head from integrin by α-actinin overexpression disrupts the establishment of integrin inter-cluster spacing.

In parallel, our data also elucidates that integrin β3, and not β1, is involved in the regulation of the inter-cluster spacing. Consistent with this, integrin β3 is involved in early signaling events by transducing mechanical signals such as small force application by magnetic beads, cyclic stretching, shear stress and substrate rigidity ^52–56^. The integrin β3 tail having opposing interactions with α-actinin and talin was also found to regulate the integrin inter-cluster spacing organization, and adhesion maturation. Competitive displacement of talin head from integrin β3 cytoplasmic tail by the integrin-binding SR12 of α-actinin disrupted organization of the integrin inter-cluster distance, which was mimicked by the integrin β3-A711P mutation (inducing a kink in the integrin β tail, leading to inactivation) that abolish the establishment of inter-cluster spacing response. Indeed, this is also consistent with previous chronological analysis of nascent adhesion formation, which documented the early recruitment of α-actinin to integrin β3^28^. This could have the effect of delaying robust activation of integrin by talin during initial adhesion engagement. Such delay could serve as a mechanical checkpoint or a coincidence detector to ensure proper engagement of key adhesome proteins and cytoskeletal connection. For example, pre-association of talin and vinculin prior to tension generation is important for subsequent adhesion maturation^57^ and proper control of rigidity sensing. Notably, aberrant expression levels of α-actinin-4 has been documented in various conditions, with elevated level linked to cancer malignancy and poor disease prognosis^58, 59^, while its loss linked to abnormal podocyte function such as in albuminuria and focal segmental glomerulosclerosis^60^. This could be due to improper spacing between integrin nanoclusters. Whether α-actinin contribute to such outcomes via its abrogation of establishment of proper inter-cluster distance and/or via talin mechanosignaling remains to be investigated in future studies, where it remains to be tested if these disease conditions are at least partially rectified by restoring proper spacing between integrin nanoclusters. Approaches described here could be used to study these diseases further.

The underlying mechanism by which cells organize nanocluster spacing and respond to inter-cluster spacing appears to be dependent on FHOD1-mediated local actin polymerization and actomyosin contractility, whereby talin head domain recruits FHOD1 to integrin nanoclusters^3^. Previous studies have also implicated FHOD1 in rigidity sensing by formation of local contractile units of actomyosin where FHOD1 can generate anti-parallel F-actin that support myosin contractility, between neighboring nanopillars spaced at ∼500 nm that potentially have integrin nanoclusters on them^18,20,61^. We postulate that FHOD1-containing mechanosensing complexes which mediate adhesion maturation on rigid substrates^31^ may thus also contribute to organization and establishment of the inter-cluster spacing described in this study. In this scenario, the optimal mechanoresponse observed when the inter-cluster spacing is in the 500-600 nm range therefore likely correspond to the size scale of the activated integrin nanocluster-talin-FHOD1-actomyosin mechanosensing complex. Due to technical and imaging challenges, visualization of the molecular players in between the nanopatterns was not possible, but can be performed in future.

The integrin-talin-FHOD1-actomyosin mechanosensing complex might act as the unifying molecular assembly that mediates not only cell sensitivity to rigidity but also to nanoscale spatial heterogeneity in ligand presentation in the microenvironments. As tissue ECM typically is assembled from biomolecules measuring in the 10-100 nm size scale^62–64^, the nanoscale ligand presentation, especially in fibrous 3D environment, is likely an intrinsic component of the cell-ECM interaction landscape^62^. Due to the continual remodeling of native ECM, an optimal inter-cluster spacing may allow cells to impose optimal ECM organization that ensure proper mechanotransduction. Conversely, loss of the mechanosensing complex in transformed cancer cells^52^, or ECM disruption due to excess deposition in fibrosis^65^, or aberrated degradation in cancer, aging, or inflammatory responses^66–69^, may in turn disrupt proper mechanotransduction response.

In conclusion, our current findings uncover an important, yet unknown, biodesign principle of self-organization in transmembrane receptor nanoclusters that is necessary for downstream signaling. In fibroblasts, for effective mechanosignal transduction, integrin receptors nanoclusters have to be spaced with a precise intercluster distance between them, and without this spatial organization, mechanosignal transduction can’t proceed. Similar transmembrane receptor spacing is observed in membrane receptors such as cadherins^4^, T-cell receptors^5,6^, receptor tyrosine kinases^8^, and this biodesign principle could be a more universal across various transmembrane receptors on the plasma membrane required for various signaling throughout the body during growth and development.

## METHODS

### Nanopattern fabrication and functionalization

Nanopatterned Ti on glass substrates were custom-ordered from ThunderNIL srl (Italy). Ti nanopatterns were functionalized by cyclo-RGD (Peptides International, cat. no. PCI-3895-PI), while glass surface was passivated by supported lipid bilayer following a previously published protocol^21^. Briefly, the Ti nanopatterned coverslips were first sonicated in 50% isopropanol. Ti was then converted to TiO_2_ by oxygen plasma (ProCleaner Plus, SKU-1062) and incubated with 1-oleoyl-2-[6-biotinyl(aminohexanoyl)]-sn-glycero-3-phosphoinositol-3,5-bisphosphate-ammonium salt (biotin PI(3,5)P_2_; 200 μg/ml for 20 min) (Avanti Polar Lipids, cat. no. 860565). The glass surface was passivated using 1,2-dioleoyl-sn-glycero-3-phosphocholine (DOPC) Small Unilamellar Vesicles (SUVs) and blocked with 1% casein for 45 min. Finally, the nanopatterns were functionalized by Dylight 650-Neutravidin (Thermofisher Scientific; cat. no. 84607; 10 mg/mL for 20 min), and then by biotin cyclo-RGD (10 mg ml^-1^ for 30 min).

### Supported lipid bilayer assembly and functionalization

Supported lipid bilayers were assembled using the SUV solution of DOPC (Avanti Polar Lipids; cat no. 850375) doped with 2% SUV solution of 1,2-dipalmitoyl-sn-glycero-3-phosphoethanolamine-N-cap biotinyl (16:0 Biotinyl Cap PE) (Avanti Polar Lipids; cat no. 870277) on acid-cleaned coverslips. The lipids were then blocked using 1-2% bovine serum albumin (Sigma-Aldrich; cat. no. A9418) for 45 mins. Subsequently, Dylight650-labeled Neutravidin (10 mg/ml for 20 min at room temperature) was used as a linker to functionalize the lipids with biotin cyclo-RGD (10 mg/ml for 30 min at room temperature).

### Glass surface functionalization

All experiments on glass substrates were performed using fibronection-coated Iwaki glass base dishes. A solution of 10 μg/mL fibronectin (BioReagent) diluted in PBS was incubated on the dishes for 1 h at 37 °C and washed thrice with PBS before cell seeding.

### Cell culture

Mouse Embryonic Fibroblasts (MEFs) ^3, 11, 53^ and Chinese Hamster Ovary (CHO.K1) cells^11^ (with low integrin expression^46^) were used for these experiments. Cells were cultured in high glucose Dulbecco’s modified Eagle’s medium (DMEM) with 10% fetal bovine serum, 1% sodium pyruvate, and 1% penicillin-streptomycin in a humidified 37°C and 5% CO_2_ incubator. Plasmids were transfected by electroporation (Neon transfection system, Life Technologies) according to manufacturer’s protocols. Experiments were carried out 42-48 h post-transfection for integrins to allow for integrin expression on the plasma membrane and 16-24 h for all other plasmids. Thirty minutes before the experiments, cells were resuspended using TrypLE express and allowed to recover in Ringer’s solution^70^. Cells were allowed to spread on the Ti nanopatterned substrates in Ringer’s solution for 1 h before fixing. MEFs were spread on fibronectin-coated glass dishes for 15 min or 1 h, as described.

### Expression Vectors

EGFP-SR12 and mCherry-Myosin-2A-N93K were a gift from Alexander Bershadsky (Mechanobiology Institute, Singapore). EGFP-Integrin β3-WT and EGFP-Integrin β3 A711P were provided by Jieqing Zhu (Versiti Inc.) ^45^. EGFP-YAP was a gift from the Boon Chuan Low (Mechanobiology Institute, Singapore). mCherry-α-actinin-4 was cloned in-house from the EGFP-ACTN4^71^ using the mCherry-N1 plasmid. mTagBFP-SR12 and mCherry-SR12 were cloned in-house from EGFP-SR12 using mTagBFP-N1 and mCherry-N1 plasmids, respectively. mTagBFP-N1, mCherry-N1, EGFP-Paxillin, mCherry-integrin αv, mCherry-Talin, EGFP-Talin head, RFP-Talin rod, and mCherry-α-actinin 1 were gifts from Michael W. Davidson (The Florida State University). mEos2-Integrin β3, Integrin β3-shRNA, tdtomato-FHOD1 and FHOD1-shRNA plasmids have been described previously^3^. Talin head-K272A, K274Q and Talin head-K322A, K324A double mutants were generated using point mutagenesis in the EGFP-Talin head plasmid.

### Antibodies

Integrin β1 blocking antibody, HMβ1-1 clone, was purchased from Biolegend (cat. no. 102202; 10 μg/mL). Phosphorylated FAK antibody was purchased from Abcam (No. ab81298; 1:200). Alexa 647-labeled secondary rabbit antibody was purchased from Invitrogen. Phalloidin-CF405M was purchased from Biotium (Cat. No. 00034; 1:200) and Phalloidin-Atto488 was purchased from Bioreagent (Cat. No. 49409; 1:200).

### Immunofluorescence

Cells were fixed with freshly prepared 4% formaldehyde (Electron Microscopy Sciences) in PBS for 15 min, permeabilized with 0.1% Triton X-100 in PBS for 20 min at 37 °C, followed by blocking with 1% BSA (Sigma-Aldrich) for 1 h at room temperature. Fixed cells were first incubated with primary antibody overnight at 4°C, followed by secondary antibody incubation with or without Phalloidin for 2 h at room temperature. Between each steps cells were washed thrice with PBS for 10 minutes each on a rocker.

### Treatment with peptides or inhibitors

For blocking integrin β3, 0.5 mM cyclic GPenRGDSPCA (GPen) peptide (Bachem) diluted in MilliQ water was used (a concentration which inhibited binding of fibronectin to αvβ3 but not to α5β1) ^72^. The cells were treated with the peptide or antibody during the 1 h spreading time.

MEFs were treated for 1 h while spreading on the nanopatterned substrates, with no pretreatment. Cells were treated with 5 μm Blebbistatin in DMSO (Sigma-Aldrich; Cat. No. B0560) for 1 h during spreading on the substrates.

### Microscopy

For single-molecule localization super-resolution imaging, a TIRF microscope (Nikon, Ti-E) equipped with a back-illuminated Scientific CMOS camera (sCMOS; Andor, Sona 4.2B-11) equipped with a 100× oil-immersion objective (Nikon CFI SR Apochromat TIRF 100× oil, 1.49 NA) was used. PALM imaging of mEos2-Integrin β3 was performed in PBS. Low intensity illumination at 405 nm was used to activate the fluorophores and 561 nm laser was used to image (and bleach) the activated fluorophores. Imaging was performed at 100 ms exposure for 7,000-10,000 frames. dSTORM imaging was performed in a reducing dSTORM buffer^73^ (50 mM Tris, 10 mM NaCl, 10 mM MEA, 0.5 mg/mL Glucose oxide, 0.1 mM catalase and 10% glucose in dH_2_O) for 20,000 frames at an exposure time of 50 ms per frame at a laser power of ∼2 kW/cm^2^. Single-molecule localizations were performed by Picasso: Localize software using a Gaussian fit mode^35^. The fitted spots were then reconstructed in Picasso, with further analysis was performed using custom MATLAB scripts.

Confocal imaging was performed using a spinning-disc confocal microscope based on a CSU-W1 Spinning Disk (Yokogawa) with a single, 70-μm pinhole disk and quad dichroic mirror along with an electron-multiplying charge-coupled device (EMCCD) (Princeton Instruments, ProEM HS 1024BX3 megapixels with 30-MHz cascade and eXcelon3 coating) using a Plan Apo 100× NA 1.45 objective lens. Super-resolution confocal imaging was performed on the same system using a LiveSR module (Roper Scientific France) which provides a structured illumination re-scanner that enhance spatial resolution ∼2X by optical deconvolution.

### Quantitative Image Processing

To calculate the period between nascent adhesions, super-resolution PALM images were used. Super-resolution images were reconstructed using Gaussian representation in Picasso software^35^. The individual localizations were exported to MATLAB where cluster analysis using DBSCAN was performed to detect individual clusters. The centers of these clusters were then analyzed using the nearest point search function in MATLAB (dsearchn) to measure the periodicity between clusters. An ideal method to measure this would be Fourier Transformation such as FFT functions. However, because focal adhesions are small and discontinuously organized on the plasma membrane, measuring the intercluster distance within the focal adhesions is not possible using FFT as FFT requires large continuous patterns. To circumvent this problem, we use the nearest point search function to handle the edge effect which arises from clusters at the very edge of focal adhesions. This algorithm measures the distance to 1 nearest neighbor and hence clusters on the edge will have the nearest cluster within the focal adhesion and it circumvents the requirement of large patterns for analysis of nearest neighbor.

Actin images were used for binary mask generation for cell area measurement^11^. Paxillin or FAK images were used to generate binary mask to measure focal adhesion area and focal adhesion length^11^. Lamellipodia was defined as the outer 5 μm area from the cell edge^79^. The percentage lamellipodia area with adhesions was measured as the total adhesion area in lamellipodia divided by total lamellipodia area. For average adhesion area per cell, all adhesions of sizes >0.1 μm^2^ were averaged for each cell. For average adhesion length per cell, all adhesions of lengths >0.3 μm were averaged for each cell. For total adhesion area per cell area, all adhesion areas were added and divided by respective cell area.

YAP nuclear to cytoplasmic ratio was assessed by measuring the intensity of a 10x10 pixel square region inside the nucleus (in the plane of maximum nuclear area) and a same-sized region in the cytosol immediately adjacent to the nuclear region. The nuclear region was defined using Hoechst 33258 (Sigma-Aldrich; cat. no. 94403).

The structural similarity was calculated using r.m.s.d. and TM-score using the TM-align tool. The AlphaFold structures^75^ of the SR12 domains in α-actinins 1 and 4 were extracted using PyMOL and aligned using TM-align^76^ (http://zhanglab.dcmb.med.umich.edu/TM-align/).

### Statistical analyses and reproducibility

All experiments were repeated between two and three times where we measured >10 cells in each repeat. Each experiment contained all the nanopatterned substrates on the same coverslip in technical duplicates, and hence all other conditions were similar, and the only variable was substrate geometry. Box plots represent interquartile distances and median, the whiskers represent the standard deviation, and the hollow circle represents the mean. Individual data points were plotted along with the box plots with *P*-values written on top of the graph.

Data from individual groups was first analyzed for normal and lognormal distribution using the Shapiro-Wilk normality test. If all groups were normally distributed, a two-tailed t-test was used for 2 groups, and ordinary one-way analysis of variance (ANOVA), followed by the Tukey’s post hoc test was used for more than 2 groups. If all groups were lognormal, the data was transformed to a normal distribution by taking the log and then analyzed. If all groups were neither normal nor lognormal, Mann-Whitney test was used for 2 groups, and Kruskal-Wallis test, followed by Dunn’s post hoc test was used for more than 2 groups. The specific test used for each graph is specified in the individual figure captions and in Supplementary Data.

## SUPPLEMENTARY INFORMATION

Supplementary Information is attached with the manuscript file.

## ACKNOWLEDGMENTS

We thank ThunderNIL Srl for fabricating the nanopatterned substrates. We thank G. Grenci (Nano and Microfabrication Core, MBI, Singapore) for useful discussions for cleaning nanopatterned coverslips, D. Pitta de Araujo (Science Communication Core, MBI, Singapore) for helpful suggestions for illustrations, and P. Kathirvel (Protein Cloning Expression Core, MBI, Singapore) for help with DNA plasmid vector cloning. We thank A. Bershadsky (MBI, Singapore), Jieqing Zhu (Versiti Inc.), B. C. Low (MBI, Singapore), and M. W. Davidson (The Florida State University, Tallahassee, FL, USA), for DNA constructs. This work was supported by intramural funds from the Mechanobiology Institute. K.J. was supported by Mechanobiology Institute Graduate Scholarship. M.P.S. received National Institutes of Health (NIH) grant support related to this project (no. RO1-GM113022). K.J. and P.K. acknowledge funding support from Ministry of Education Academic Research Fund Tier2 (MOE2019-T2-2-014).

## AUTHOR CONTRIBUTIONS

K.J. & R.C. designed experiments with help from M.P.S. and P.K. K.J. performed the experiments. R.C. has established protocols for Ti functionalization, has cloned several of the mutant constructs used here and helped establish data analysis protocols. K.J. analyzed the data. K.L. provided assistance in carrying out experiments on glass dishes and their analysis. K.J., R.C., M.P.S. and P.K. wrote the manuscript.

