## Supplementary Information for "Intrinsic self-organization of integrin nanoclusters within focal adhesions is required for cellular mechanotransduction"

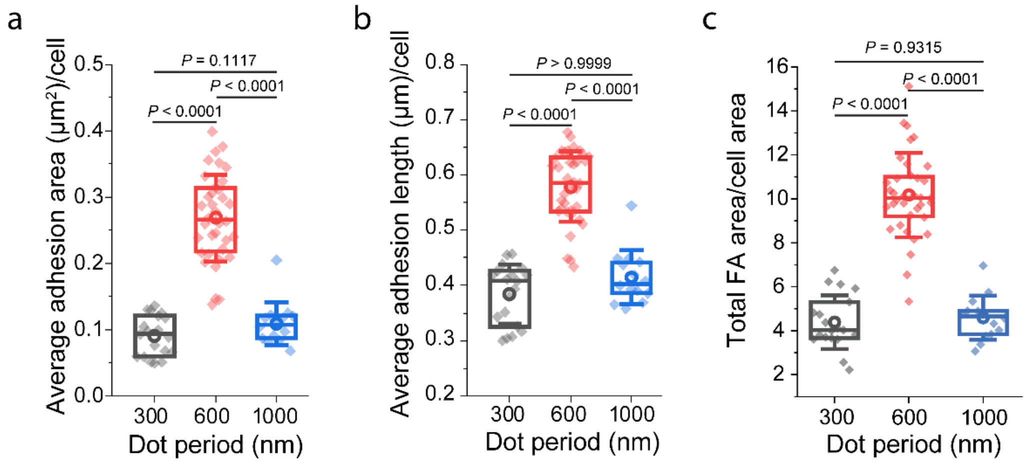


Figure S1. *MEFs show an increased adhesion area and adhesion length on intercluster spacing of 600 nm.*

a) average adhesion area (μm$^2$) per cell for the different dot periods (n ≥ 14 cells from three independent experiments, analyzed by the one-way ANOVA test and Tukey's post hoc test; P values are indicated in the figure). b) average adhesion length (μm) per cell for the different dot periods (n ≥ 14 cells from three independent experiments, analyzed by the Kruskal–Wallis test and Dunn's post hoc test; P values are indicated in the figure).c) percentage of cell area with adhesions calculated as total FA area per cell area (n ≥ 14 cells from three independent experiments, analysed by the One-way ANOVA test and Tukey's post hoc test; P values are indicated in the figure). Box plots with data overlay display the upper and lower quartiles and a median, the circle represents the mean, and the whiskers denote the standard deviation values, along with the individual points.

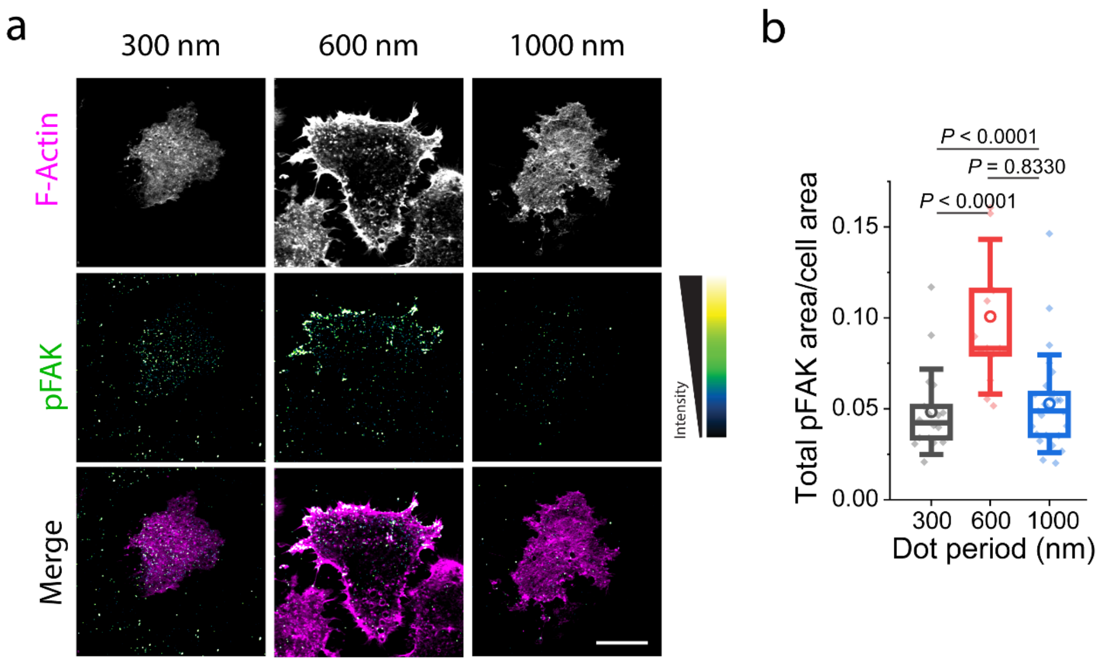


*<u>Fig. S2.</u> MEFs show an increased phosphorylation of FAK on intercluster spacing of 600 nm.*
a, Representative fluorescence images of F-actin and phospho-FAK stained MEFs, along with the merge, on the different disc periods. b, Quantifications of Total FAK area/cell area (n≥14 cells from three independent experiments, analysed by one-way ANOVA and Tukey's post hoc test). Box plots with data overlay display the upper and lower quartiles and a median, the circle represents the mean, and the whiskers denote the standard deviation values, along with the individual points.

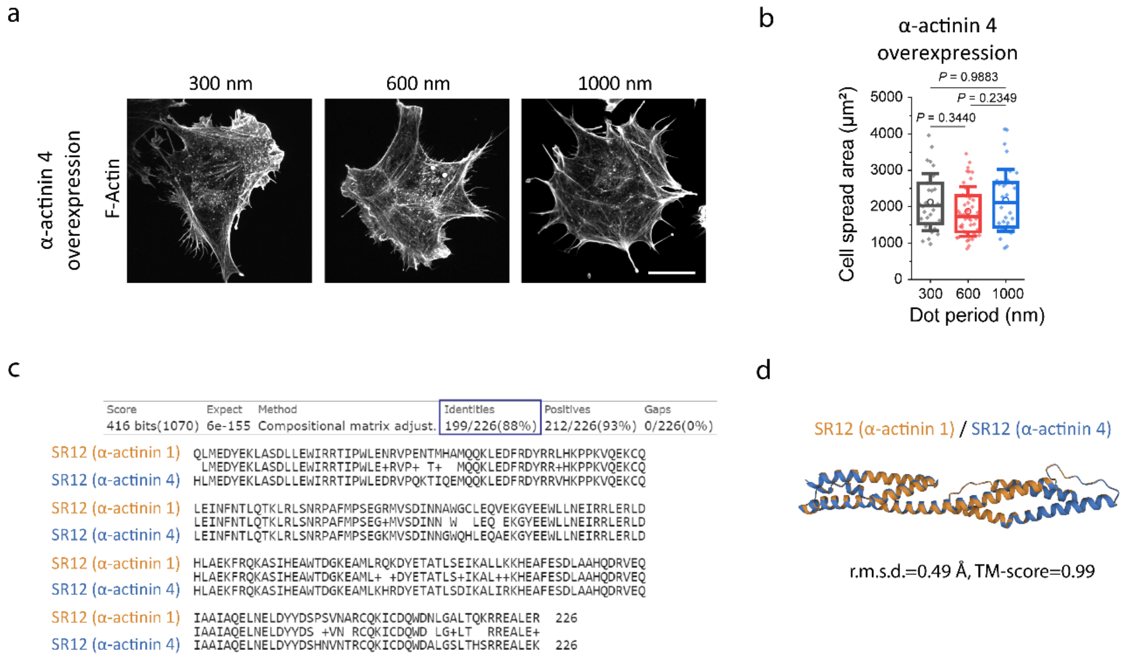


*<u>Fig. S3.</u> Overexpression of the α-actinin 4 isoform phenocopies α-actinin 4 and disrupts integrin intercluster spacing-dependent cell spreading.*

a, Representative F-actin images of cells overexpressing mCherry-α-actinin 4 on the different disc periods, Scale bar, 20 μm, along with b, quantification of the cell spread area (n≥30 cells from three independent experiments, analyzed by one-way ANOVA test and Tukey's post hoc test; *P* values are indicated). c, Protein sequence analysis of the SR12 domains of α-actinin 1 (orange) and α-actinin 4 (blue) using NCBI BLAST, showing 88\% identities in both the sequences. d, Structural analysis of the SR12 domains of α-actinin 1 (orange) and α-actinin 4 (blue) using the TM-align tool, showing an r.m.s.d. of 0.49 Å and a TM-score of 0.99. Box plots with data overlay display the upper and lower quartiles and a median, the circle represents the mean, and the whiskers denote the standard deviation values, along with the individual points.

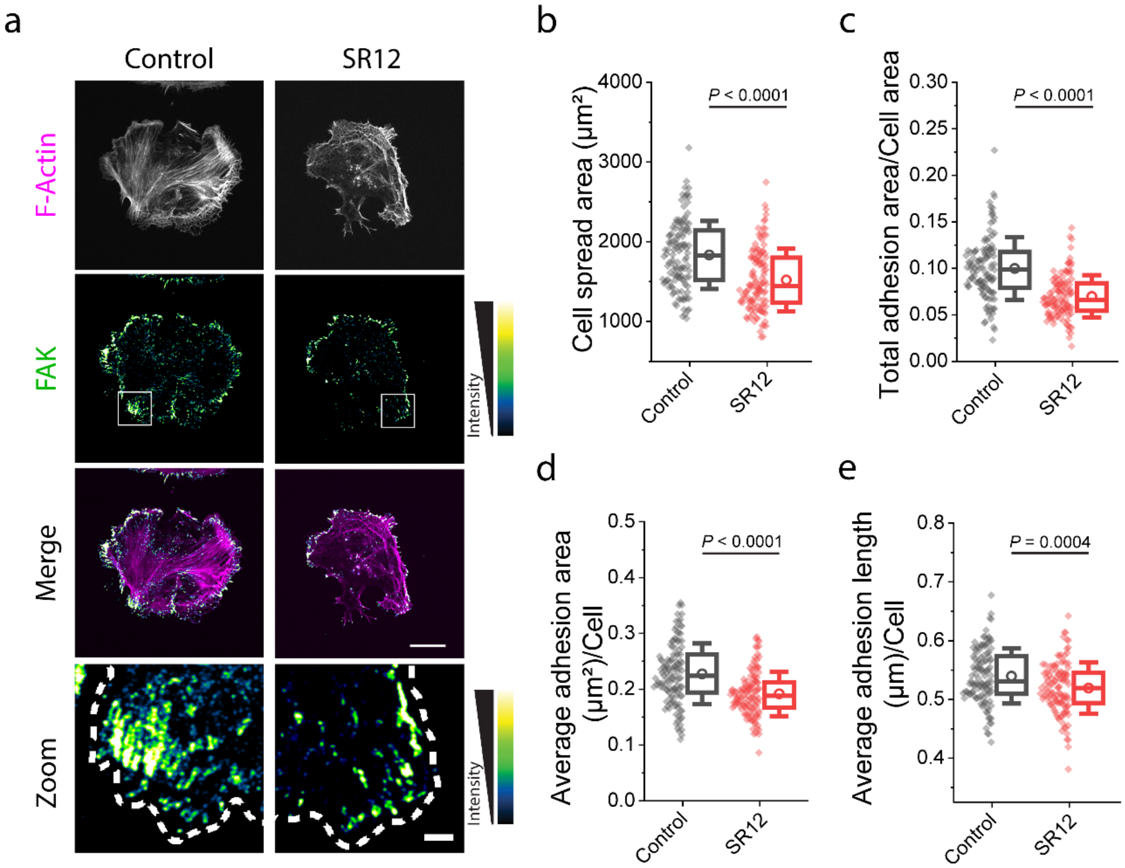


*Fig. S4. SR12 overexpression reduces cell spreading and impedes adhesion maturation in early spreading as seen by reduced adhesion density, length, and area.*

a, Control and SR12 overexpressing cells were spread on fibronectin-coated glass dishes for 15-20 mins and immunostained for FAK and actin (phalloidin). Representative images showing F-actin, FAK, merge, and zoom of the white box in FAK. Scale bar for Merge, 20 µm. Scale bar for zoom, 2 µm. b, Quantification of the cell spread area (n≥118 cells for control and 117 cells for SR12 from three independent experiments, analyzed by unpaired two-tailed t-test; *P* value is indicated), c, total adhesion area normalized to the cell area (analyzed by two-tailed Mann Whitney test; *P* value is indicated), d, average adhesion area per cell (analyzed by unpaired two-tailed t-test; *P* value is indicated), and e, average adhesion length per cell (analyzed by unpaired two-tailed t-test; *P* value is indicated). All graphs show box plots of interquartile distance, the line represents the median, the circle represents the mean and the whiskers represent the standard deviation.

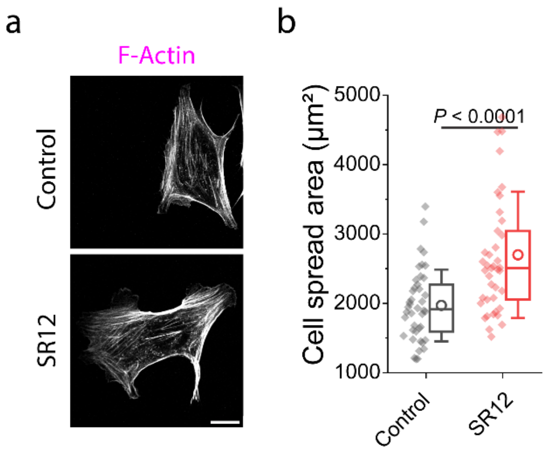


*<u>Fig. S5.</u> SR12 overexpression increases cell spreading after 1 hr spreading.*

a, Control and SR12 overexpressing cells were spread on fibronectin-coated glass dishes for 1 hr and immunostained for actin (phalloidin). Scale bar, 20 µm. b, Quantification of the cell spread area (n≥42 cells for control and 41 cells for SR12 from three independent experiments, analyzed by unpaired two-tailed t-test; *P* value is indicated). All graphs show box plots of interquartile distance, the line represents the median, the circle represents the mean and the whiskers represent the standard deviation.

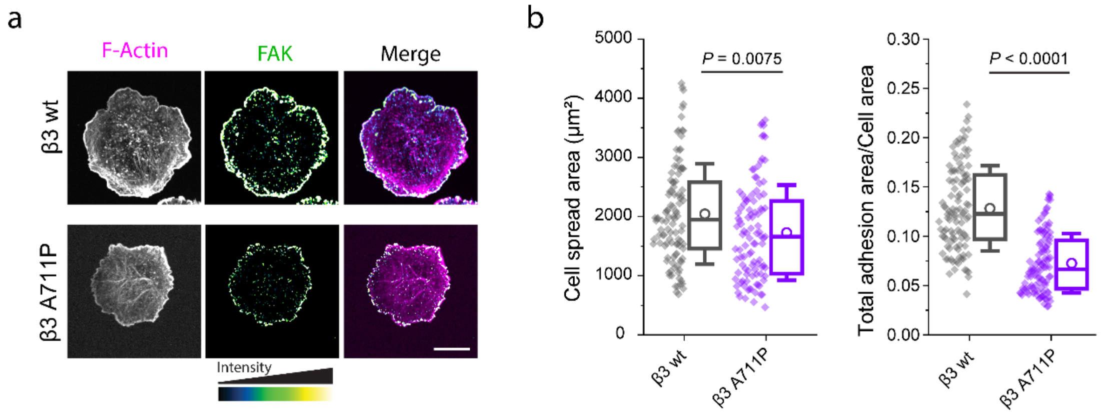


*Fig. S6. The A711P mutation on Integrin β3 impedes cell spreading and adhesion maturation.*
a, Representative phalloidin-labeled F-actin, FAK and merge images of integrin β3 wild-type and β3 A711P mutant expressing CHO cells on fibronectin-coated glass dishes for 20 mins. Scale bar, 15 µm. b, Quantification of the cell spread area (n≥103 cells from three independent experiments, analyzed by Mann-Whitney test) and total adhesion area normalized to the cell area (analyzed by Mann-Whitney test). Box plots with data overlay display the upper and lower quartiles and a median, the circle represents the mean, and the whiskers denote the standard deviation values, along with the individual points.

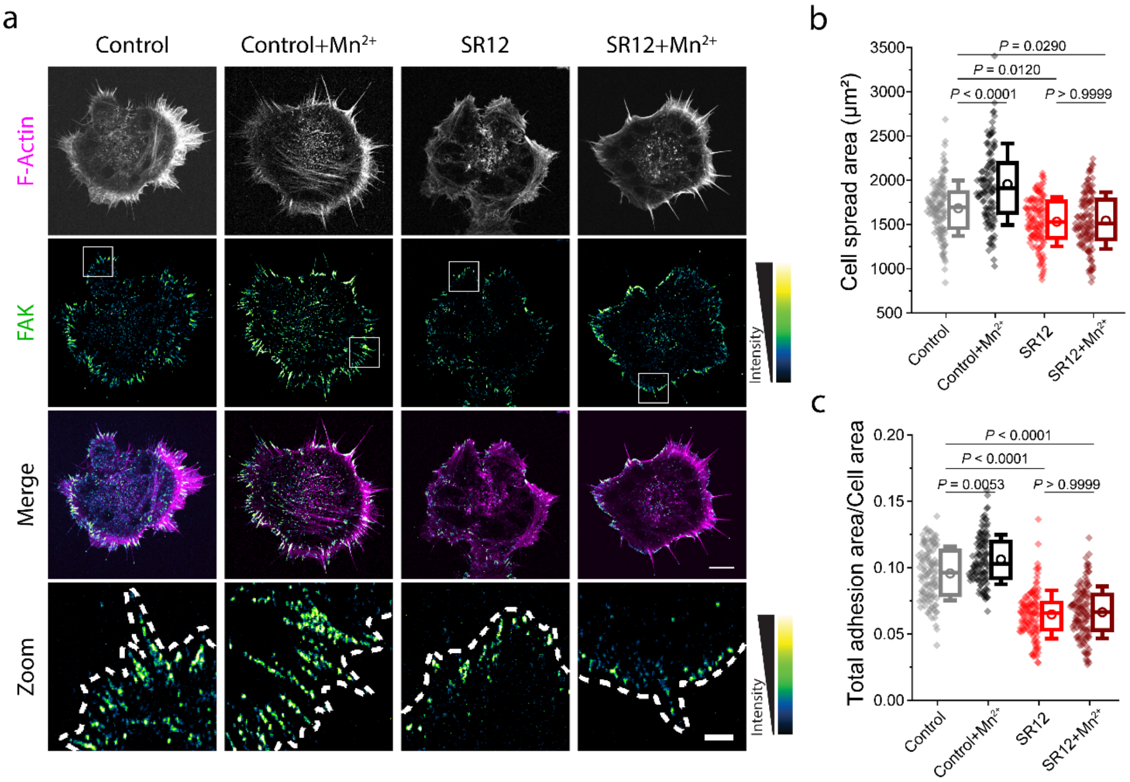


*<u>Fig. S7.</u> Integrin activation by $Mn^{2+}$ does not rescue the loss of adhesion maturation by SR12.*

a, Representative images of control cells, $Mn^{2+}$ treated control cells, SR12 overexpressing cells, and $Mn^{2+}$ treated SR12 overexpressing cells immunostained for FAK, actin (phalloidin), merge and zoom of the white box in FAK. b, Quantification of the cell spread area (n≥112 cells from three independent experiments, analyzed by Kruskal-Wallis test and Dunn's post hoc test; *P* values are indicated), and c, total adhesion area normalized to the cell area (analyzed by Kruskal-Wallis test and Dunn's post hoc test; *P* values are indicated). All graphs show violin plots and box plots of interquartile distance, the notch represents the median, the square represents the mean and the whiskers represent the standard deviation.

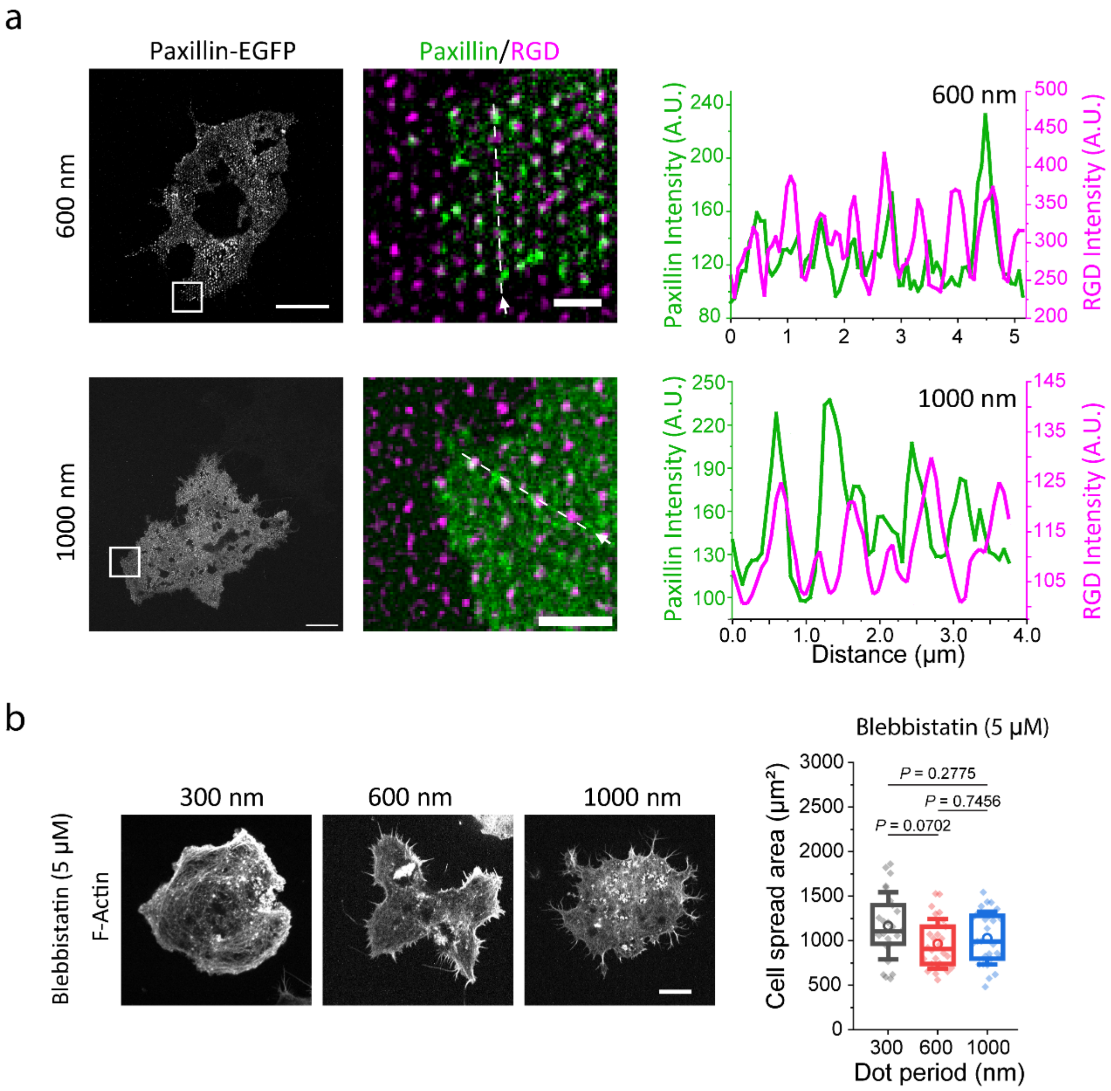


*Fig. S8. Reduced myosin activity leads to loss of establishment of intercluster spacing.*

a, SR-confocal fluorescence images of EGFP-paxillin (left) images on 600 nm nanodisc period (top) or on 1000 nm nanodisc period (bottom). A zoom of the white box along with the RGD functionalized (marked using Neutravidin) nanodisc substrates (center). Fluorescence intensity line plot of paxillin and RGD along the white dotted line. Scale bar (left), 10 µm. Scale bar (center), 1 µm. b, Representative phalloidin-labelled F-actin images of 5 µM blebbistatin treated MEFs spread on the different disc periods along with quantification of the spread area of cells spread on the different disc substrates. (n≥25 cells from three independent experiments, analysed by the one-way ANOVA and Tukey's post hoc test; *P* values are indicated in the figure). Scale bar, 10 µm. Box plots with data overlay display the upper and lower quartiles and a median, the circle represents the mean, and the whiskers denote the standard deviation values, along with the individual points.
